# Estimating the impact of pneumococcal conjugate vaccines on childhood pneumonia in sub-Saharan Africa: A systematic review

**DOI:** 10.1101/865154

**Authors:** Chukwuemeka Onwuchekwa, Edem Bassey, Victor Williams, Emmanuel Oga

## Abstract

**Background:** The impact of pneumococcal conjugate vaccine introduction in reducing the incidence of childhood pneumonia has not been well documented in sub-Saharan Africa. Many studies evaluating vaccine impact have used invasive pneumococcal disease or pneumococcal pneumonia as an outcome.

**Objective:** To estimate the impact of routine administration of 10-valent and 13-valent PCV on the incidence of pneumonia in children under five years of age in sub-Saharan Africa.

**Data sources:** A systematic review was conducted between 16 and 31 July 2019. The review was registered on PROSPERO with registration number CRD42019142369. The literature search was conducted in indexed databases including Medline and Embase, grey literature databases and online libraries of two universities. Manual search of the references of included studies was performed to identify additional relevant studies. The search strategy combined pneumococcal conjugate vaccine, pneumonia and child as search concepts.

**Study selection:** Studies investigating the impact of 10- or13-valent PCV on childhood pneumonia in a sub-Saharan African country were eligible for inclusion. Case-control, cohort, pre-post and time-series study designs were eligible for inclusion. Exclusion criteria were use of 7- or 9-valent PCV, systematic review studies, clinical trials and record publication prior to 2009.

**Data extraction:** Independent data extraction was conducted. Key variables include year study conducted, type of study design, type of PCV used and year of introduction, reported PCV coverage, outcome measure evaluated and the effect measure.

**Data synthesis:** Eight records were included in the final analysis, 6 records were pre-post or time-series studies, 1 was a case-control study and 1 report combined pre-post and case-control studies. Vaccine impact measured as percentage reduction in risk (%RR) of clinical pneumonia was mostly small and non-significant. The risk reduction was more significant and consistent on radiological and pneumococcal pneumonia. Vaccine effectiveness reported in case-control studies was mostly non-significant.

**Conclusion:** Evidence of the positive impact of routine infant pneumococcal vaccination on pneumonia in sub-Saharan Africa is weak. There is a need for more research in this area to evaluate the influence of pathogen or serotype replacement in pneumonia after PCV introduction. Ongoing surveillance is also required to establish the long term trend in pneumonia epidemiology after PCV introduction.

## Introduction

### Rationale

Pneumonia is one of the leading causes of childhood deaths globally, particularly in Africa (1). Despite consistent year-on-year reduction in the total number of childhood deaths attributed to pneumonia or lower respiratory infections, pneumonia still account for a significant proportion of all childhood deaths globally. According to the World Health Oganization (WHO) estimates, the African region accounted for about half of the 808,694 childhood deaths due to lower respiratory infections. Annually over a 100 million cases of pneumonia are reported in children less than five years of age, mostly in poor countries in Africa and Asia(2–4).

Streptococcus pneumonia is the major cause of childhood pneumonia deaths and is the leading cause of vaccine-preventable child deaths globally(5–7). Pneumococcal pneumonia causes between 1 and 4 million episodes of pneumonia in Africa yearly (8). *Streptococcus pneumonia* is a gram-positive diplococcic that usually colonizes the human upper respiratory tract and can occasionally be found in the airways of other animals(9–14). Infection occurs mostly through autoinfection when the bacteria is translocate to the normally sterile environment of the lower tract and evades host immune defences (9,12,14,15). There are 97 serotypes of *S. pneumoniae* currently known based on their polysaccharide capsule antigen(16). The 97 serotypes differ in carriage potential and propensity to cause invasive disease including pneumonia, otitis media and meningitis (17,18), with the thirteen most common serotypes accounting for 70 – 75% of invasive pneumococcal disease globally(8,18,19).

Pneumococcal conjugate vaccines (PCV) have been licensed since 2000; previous polysaccharide-based vaccines were found to have poor immunogenicity in children(8). An initial 7-valent PCV included serotypes 4, 6B, 9V, 14, 18C, 19F and 23F providing cover against 67% of disease-causing serotypes. The 10-valent and 13-valent PCV include in addition to the 7-valent serotypes 1, 3 and 7F; and 1, 3, 7F, 19A, 6A and 3. The 10-valent and 13-valent vaccine serotypes cover over 80% of invasive disease-causing serotypes(8,20,21). Sub-Saharan African countries like South-Africa and The Gambia introduced the 7-valent PCV into routine infant vaccination schedule in the later part of the last decade(22,23). The WHO recommends the 10- or 13-valent PCV for routine infant immunization. Many countries in Africa through the support of Global Alliance for Vaccination and Immunization (GAVI) have incorporated pneumococcal vaccines into their expanded program of Immunization (EPI) schedules. With vaccine coverage above 90% in most countries in sub-Saharan Africa, PCV vaccination is associated with a significant decline in invasive pneumococcal disease incidence globally and at individual country level (16,22,23). The benefit of vaccines post introduction is usually estimated by measuring vaccine effectiveness or vaccine impact. Vaccine effectiveness measures the reduction in disease rates in the context of routine real world use; this is usually done in the context of case-control or Cohort studies. Vaccine impact on the other hand quantifies the reduction in disease rate at the population level following vaccine introduction. Vaccine impact is usually measured by quasi-randomised methods like pre-post studies and interrupted time series analysis(24)

The impact of vaccination on pneumococcal pneumonia has been clearly demonstrated; however its impact on clinical and radiological pneumonia has been more modest(23). Studies in sub-Saharan Africa have reported varied measures of vaccine effectiveness and impact. Overall, there is overwhelming evidence of effectiveness against invasive pneumococcal disease with significant decline in disease after introduction of the vaccine (25–29).

### Objectives

This review seeks to evaluate the impact of routine administration of pneumococcal conjugate vaccines on clinical pneumonia, radiological pneumonia and pneumococcal pneumonia in children under five years of age in sub-Saharan Africa. The research objective was formulated using the PICO (population, intervention, comparison and outcome) framework and summarised in Table 1(32)

**Table 1.** PICO framework for formulating the review question.

| Concept | Elaboration |
| --- | --- |
| Population of interest | Children between 1 – 59 months of age from any of the 46 countries in sub-Saharan Africa. |
| Intervention | Routine immunization with 10- or 13-Valent PCV |
| Comparison (or control) | Populations before and after PCV introduction |
|  | OR<br>Comparison between groups with (cases) and without (controls) the outcomes of interest |
| Outcomes | Clinical pneumonia, radiological pneumonia and pneumococcal studies |

## Methods

### Study protocol

The systematic review protocol was developed in line with PRISMA reporting guidelines and available as supplementary material (33). The review was registered on PROSPERO with registration number CRD42019142369.

### Eligibility criteria

We included primary individual and population-based studies conducted in sub-Saharan Africa evaluating 10-valent or 13-valent PCV impact in children published since 1 January 2010. This time was chosen because the earliest countries in Africa introduced PCV into routine infant vaccine schedules in 2009. Studies conducted in children between 1 and 59 months of age within any sub-Saharan African location were eligible for inclusion.

Studies that had pneumonia as an outcome were eligible for inclusion, this included clinical pneumonia, radiological pneumonia, pneumonia-related hospitalizations and pneumonia-related deaths. Additionally, studies that applied invasive pneumococcal disease (IPD) as outcome were also considered for inclusion if pneumonia was reported separately.

Study types eligible for inclusion were pre-post quasi-experimental designs, interrupted time series, Cohort and Case-control studies. For pre-post studies and interrupted time series, we included only studies where the outcome was assessed 3 years or more after vaccine introduction.

Studies using 7-valent or 9-valent PCV, review papers, clinical trials and papers without available versions in English were excluded. We excluded 7-valent PCV studies because these vaccines were shown to have limted serotype coverage in sub-Saharan Africa.

### Information sources

We conducted a search of peer-reviewed and grey literature relating to the study question. MEDLINE search was conducted in PubMed on 16 Jul 2019 with final search on 31 Jul 2019. Scopus search was conducted on 20 Jul 2019, Embase (Ovid) was searched on the 29 Jul 2019. The search also included Africa-wide information (29 Jul 2019) and African index Medicus (24 Jul 2019). Grey literature searches included Open grey (20 Jul 2019), ProQuest Dissertation & theses global (20 Jul 2019), London school of hygiene and tropical medicine research online (22 Jul 2019) and University of Edinburgh library (28 Jul 2019).

### Search

The search strategy combined the key concepts of the research question and based on the PICO framework. The three components or concepts were: population of interest (children below 5 years of age), intervention being investigated (pneumococcal conjugate vaccine) and the outcome of interest (pneumonia). The boolean operators AND and OR were used to combine the search concepts.

Further details of the database search strategy and date of searches can be found in supplementary materials

### Study selection process

Two members of the review team conducted the database screening independently. Reading through the titles and abstracts of the search results we identified records to be included in the full-text screening process based on the eligibility criteria. Records for which there was uncertainty or disagreement about eligibility during the title and abstract screening were included for full-text screening. The second stage of the screening involved reading the full-text of the records to determine if they were eligible for inclusion. Finally, a process of backward-snowballing was performed on all eligible papers. Here we searched through the references of eligible papers for other relevant publications that could be included in the review. The PRISMA flow diagram for the study selection procedure is shown in Figure 1.

**Figure 1.**
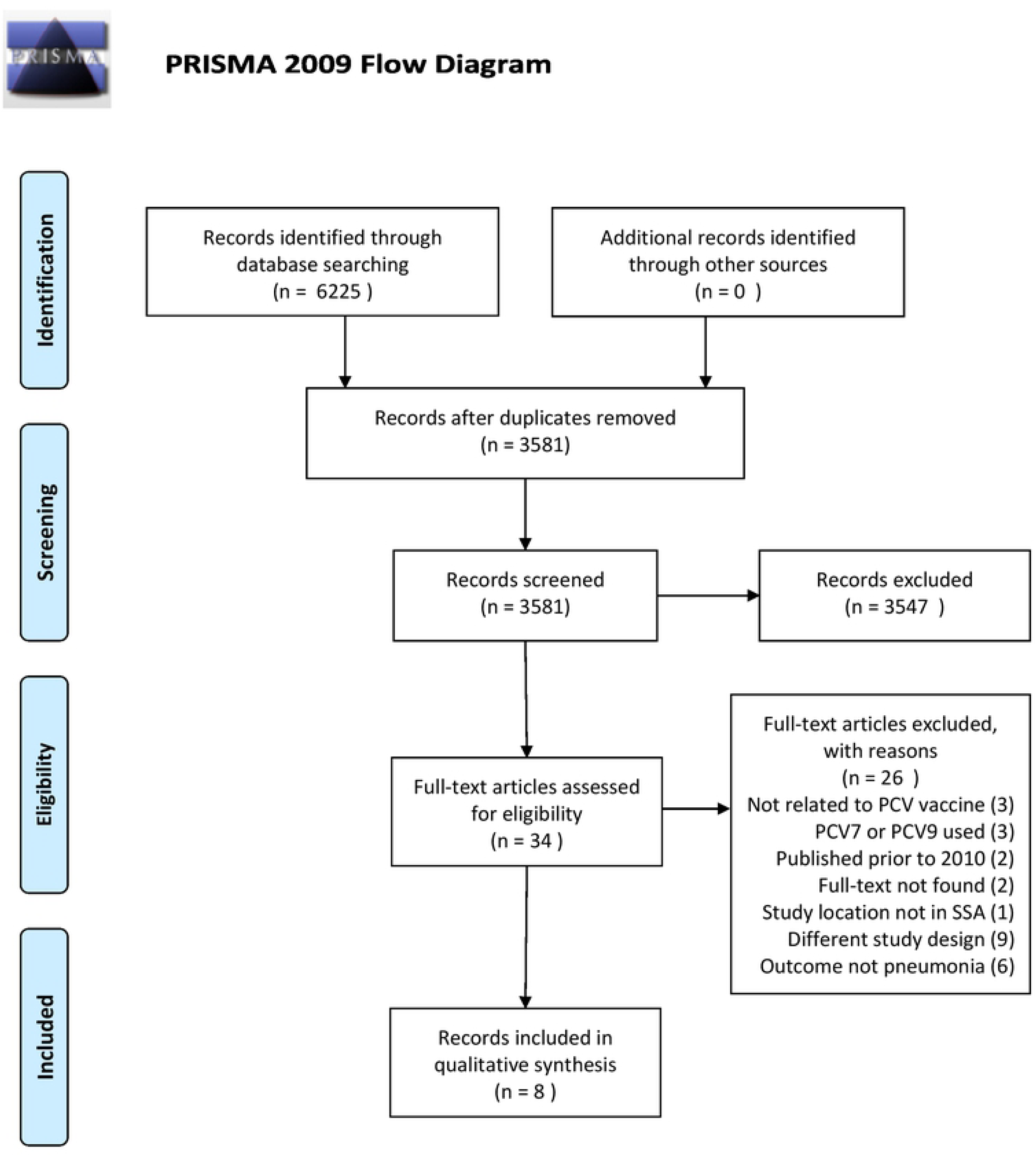

All stages of the publication search were managed using a citation management application (Mendeley).

### Data collection process and Data items

The following information was collected for each included record: first author name, year of publication, Publication type (full-text or abstract), study location and country, location type (hospital-based or population-based), study design, study aim (to assess impact or effectiveness), data sources (clinical or laboratory), study population description, HIV status of participants, mean or median age, sex distribution, type of PCV in current use (PCV 10 or PCV13), year PCV7 introduced, year PCV10 or PCV13 introduced, minimum reported coverage during PCV10 or PCV13 period, Baseline and post-vaccination periods (for pre-post and interrupted time-series), outcome measure, outcome definition, outcome measure and confidence interval if reported. The information was collected directly into an extraction form in Excel by one member of the review team and confirmed by another. No additional information was sought from investigators or authors.

### Risk of bias in individual studies

We assessed the quality of each study included in the review. For the case-control studies assessing vaccine effectiveness, we used the National Heart, Lung and Blood Institute Quality assessment tool for case-control studies. We adapted the National Heart, Lung and Blood institute Quality Assessment Tool for Before-After (Pre-Post) Studies With No Control Group for pre-post and interrupted time series analyses(34). The study quality assessment tools are presented as supplementary material.

### Summary measures

The primary outcomes of interest in this review were clinical pneumonia and radiological pneumonia. The secondary outcome was pneumococcal pneumonia. There was variation in the ascertainment of the outcomes and outcome measures between studies. In pre-post and interrupted time-series studies comparing outcome incidence before and after PCV introduction, measures were presented as percentage reduction in incidence and incidence ratios. Where possible incidence ratios were converted to percentage reduction in incidence: *% reduction in risk = (1 – aRR) X 100%;* where aRR is the adjusted Risk/Rate ratio for post- and pre-vaccination periods(35). In case-control studies we presented adjusted vaccine effectiveness (aVE) as reported by the authors. When adjusted odd Ratios (aOR) were presented we estimated aVE as: *Adjusted vaccine effectiveness = (1 – aOR) X 100%* (35); where aOR = adjusted odd ratio.

### Synthesis of results

Due to the variability in study designs, recruitment method and definition of pneumonia applied, we present a narrative synthesis of the included studies for the impact of PCV on childhood clinical pneumonia, radiological pneumonia and pneumococcal pneumonia.

## Results

### Study selection

In total eight records reporting on nine studies were included in the review. The literature search produced 3581 distinct records after removing duplicate records. After screening the titles and abstracts, we identified 34 relevant records for the full-text review. We excluded a further 26 records after full-text assessment. Three records were unrelated to PCV introduction, one record was from outside sub-Saharan Africa and three record reported on 7- or 9-valent PCV. Two records had studies conducted prior to 2010, nine records had ineligible study designs and six records had ineligible outcome (including invasive pneumococcal disease). We could not retrieve full-text titles for two records. Backward snowballing of the included records produced no additional records for inclusion in the review. Figure 1 is a flow diagram depicting the record selection process.

### Study characteristics

Of the eight records included in the review; four were conducted in South Africa, two were conducted in Kenya, and one each from Rwanda and The Gambia. There were 7 studies with pre-post or interrupted time-series design and two case-control studies (Mackenzie et al included report on a case-control study). Most of the study locations had 7-valent PCV introduced before transition to the 13-valent PCV except in Kenya where the 10-valent PCVwas used without a 7-valent PCV period.

**Table 2.** Characteristics of studies included in review

| **Author** | **Country** | **Study design** | **Data sources** | **Population** | **PCV** | **Year PCV7 intro** | **Year PCV intro** | **baseline period** | **Post-intervention period** |
| --- | --- | --- | --- | --- | --- | --- | --- | --- | --- |
| Von Mollendorf et al, 2017(36) | South Africa | Pre-post | Laboratory surveillance | < 60 months | 13 | 2009 | 2011 | 2005 – 2008 | 2012 - 2013 |
| Mackenzie et al, 2017(37) | URR, The Gambia | Pre-post | population surveillance | 2 to 59 months | 13 | 2009 | 2011 | 2008 – 2010 | 2014 - 2015 |
| Mackenzie et al, 2017*(37) | URR, The Gambia | case-control | Clinical | 3 to 59 months | 13 | 2009 | 2011 | NA | NA |
| Hammit et al, 2019(38) | Kilifi, Kenya | pre-post | clinical surveillance | < 60 months | 10 | NA | 2011 | 1999 – 2010 | 2012 - 2016 |
| Gatera et al, 2016(31) | Rwanda (five districts) | Pre-post | Clinical and laboratory surveillance | < 60 months | 13 | 2009 | 2011 | 2002 – 2009 | 2012 |
| Silaba et al, 2019(39) | Kilifi, Kenya | ITS | population surveillance | 2 to 59 months | 10 | NA | 2011 | 2002 – 2011 | 2011 - 2015 |
| Mahdi et al, 2015(40) | Soweto, Cape town, KwaZulu-Natal. South Africa | case-control | clinical | 2 to 42 months | 13 | 2009 | 2011 | NA | NA |
| Solomon et al, 2017(41) | Soweto, South Africa | Pre-post | population surveillance | < 60 months | 13 | 2009 | 2011 | 2006 – 2008 | 2010 – 2014 |
| Tempia et al, 2015(42) | Soweto, South | Pre-post | laboratory surveillance | < 24 months | 13 | 2009 | 2011 | 2009 - | 2011 – 2012 |

\*Reported with the pre-post study.

### Risk of bias within studies

Five of the seven population-based studies were considered to be of good quality with low risk of bias. The two case-control reports were graded as having good quality. Details on the study quality assessment are available in Appendix 5.

### Result of individual studies

Individual studies showed variability in their definition of clinical, radiological and identification of pneumococcal pneumonia. Most studies applied the WHO standards for identification of Radiological pneumonia and clinical pneumonia (23,39,43). One study based pneumonia diagnosis on ICD-10 coding from clinical notes(44). Studies reporting on pneumococcal pneumonia were based on culture with occasional confirmation by Polymerase chain reaction (PCR) (42,45,46) except in one study where PCR based method was used as a means of surveillance(42).

**Table 3.** Summary of effect reported in studies included in the review.

|  | Outcome | Clinical pneumonia | Radiological pneumonia | Pneumococcal pneumonia |
| --- | --- | --- | --- | --- |
|  |  | Percent risk reduction (95% CI) | Percent risk reduction (95% CI) | Percent risk reduction (95% CI) |
| Von Mollendorf et al, 2017(36) | < 60 months | -- | -- | 65 (64 – 67) |
|  | HIV positive | -- | -- | 51 (49 – 55) |
|  | 12 – 59 months | -- | -- | 69 (67 – 71) |
| Mackenzie et al, 2017(37) | 2 – 11 months | 2 (-4 – 8) | 23 (7 – 36) | -- |
|  | 12- 23 months | -6 (-15 – 2) | 29 (12 - 42) | -- |
|  | 24 – 59 months | -7 (-18 – 2) | 22 (1 - 39) | -- |
| Hammit et al, 2019(38) | < 60 months | -- | -- | 85 (66 – 93) |
| Gatera et al, 2016(31) | < 60 months | 15 (Not reported) | -- | -- |
| Silaba et al, 2019(39) | 2 – 59 months | 27 (3 – 46) | 48 (14 – 68) | -- |
|  | 2 – 11 months | -- | 27 (-36 – 61) | -- |
|  | 12 – 23 months | -- | 46 (-10 – 73) | -- |
|  | 24 – 59 months | -- | 11 (-69 – 53) | -- |
| Solomon et al, 2017(41)<br>** | HIV infected | 33 (6 -52)¶¶ | -- | -- |
|  | HIV uninfected or unconfirmed | 39 (24 – 50) ¶¶ | -- | -- |
|  | 3 – 11 months | 39 (10 – 57) ¶¶ | -- | -- |
|  | 12 – 23 months | 15 (-37 – 42)¶¶ | -- | -- |
|  | 24 – 59 months | 45 (17 – 62)¶¶ | -- | -- |
| Tempia et al, 2015(42) | HIV uninfected | -- | -- | 66.8 (43.8 – 81.3) |
|  |  | -- | -- | 64.0 (52.6 – 81.2) |
|  |  | **Adjusted vaccine effectiveness (95% CI)** | **Adjusted vaccine effectiveness (95% CI)** | **Adjusted vaccine effectiveness (95% CI)** |
| Mackenzie et al, 2017*(37) | 3 – 59 months | -- | 28 (-23 – 58) | -- |
| Mahdi et al, 2015(40) | Hospital controls | -- | 20.0 (-9.3 – 41.6) | -- |
|  | Community controls | -- | 32.1 (4.6 – 51.6) | -- |

*\*Based on WHO clinical classification of severe and very severe pneumonia* *\*\* pneumonia hospitalization based on ICD-10 coding.* *¶ 50% credible interval based on Bayesian methods (negative values indicate an increase in incidence).* *€ Identification based on polymerase chain reaction* *± pneumonia classified based on WHO criteria or based on abnormal CXR plus CRP >40mg/L*

## Synthesis of results

### Change in clinical pneumonia

Four population based studies evaluated the impact of 10- or 13-valent vaccine on clinical pneumonia with inconsistent findings. Silaba et al. reported a decline in hospitalization for severe or very severe pneumonia based on WHO clinical definition(47). Mackenzie et al. showed no significant decline in clinical pneumonia incidence in all under-five age groups three years after 13-valent PCV introduction. Solomon et al. reported a significant decline in pneumonia in infants and children between 24 – 59 months. This effect was also observed among HIV infected and HIV uninfected or HIV status unknown children(41). Figure 2 graphically displays the reported reduction in clinical pneumonia incidence.

**Figure 2.**
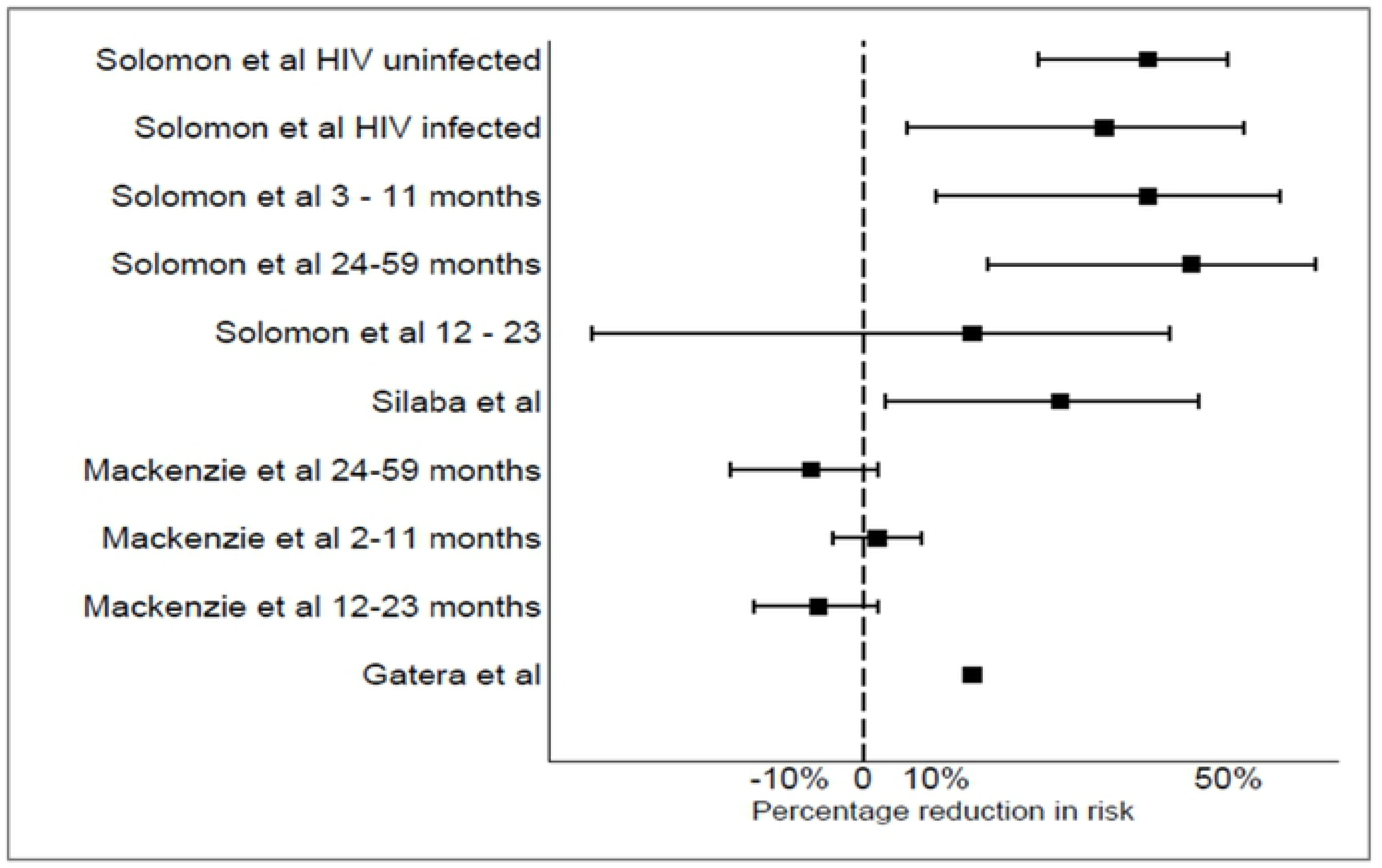
Forest plot showing the percentage risk reduction in clinical pneumonia incidence in included studies and population.

### Change in Radiological pneumonia

Two impact studies evaluated radiological pneumonia as an outcome(23,47). Mackenzie et al reported a decline in WHO defined radiological pneumonia in all age groups with decline in the range of 22 – 29%. This decline was most pronounced in the 12 – 23 month age group(23). Silaba et al. also reported a 48% decline in radiological pneumonia in the entire under-five population. A similar trend was observed in the age groups with the 12 – 23 month group experiencing the greatest reduction in radiological pneumonia(47).

The case-control study reported by Mackenzie et al. using community controls showed a vaccine effectiveness of about 28% however this did not reach statistic significance. Mahdi et al. reported adjusted vaccine effectiveness measures using both community and hospital controls. Vaccine effectiveness was significant with community controls (aVE 32.1%, 95% CI 4.6% - 51.6%). Vaccine effectiveness was however not significant with hospital controls (aVE 20%, 95% CI −9.3% – 41.6%).

### Change in Pneumococcal pneumonia

All three studies that reported on pneumococcal pneumonia were based on microbiological diagnosis with PCR confirmation. The report by Tempia et al. provided separate results based on PCR identification and mircobiological identification. Included studies consistently showed decline in cases of pneumococcal pneumonia after vaccine introduction irrespective of method of pneumococcal identification. Figure 3 shows the reduction in pneumococcal pneumonia incidence reported from included studies

**Figure 3.**
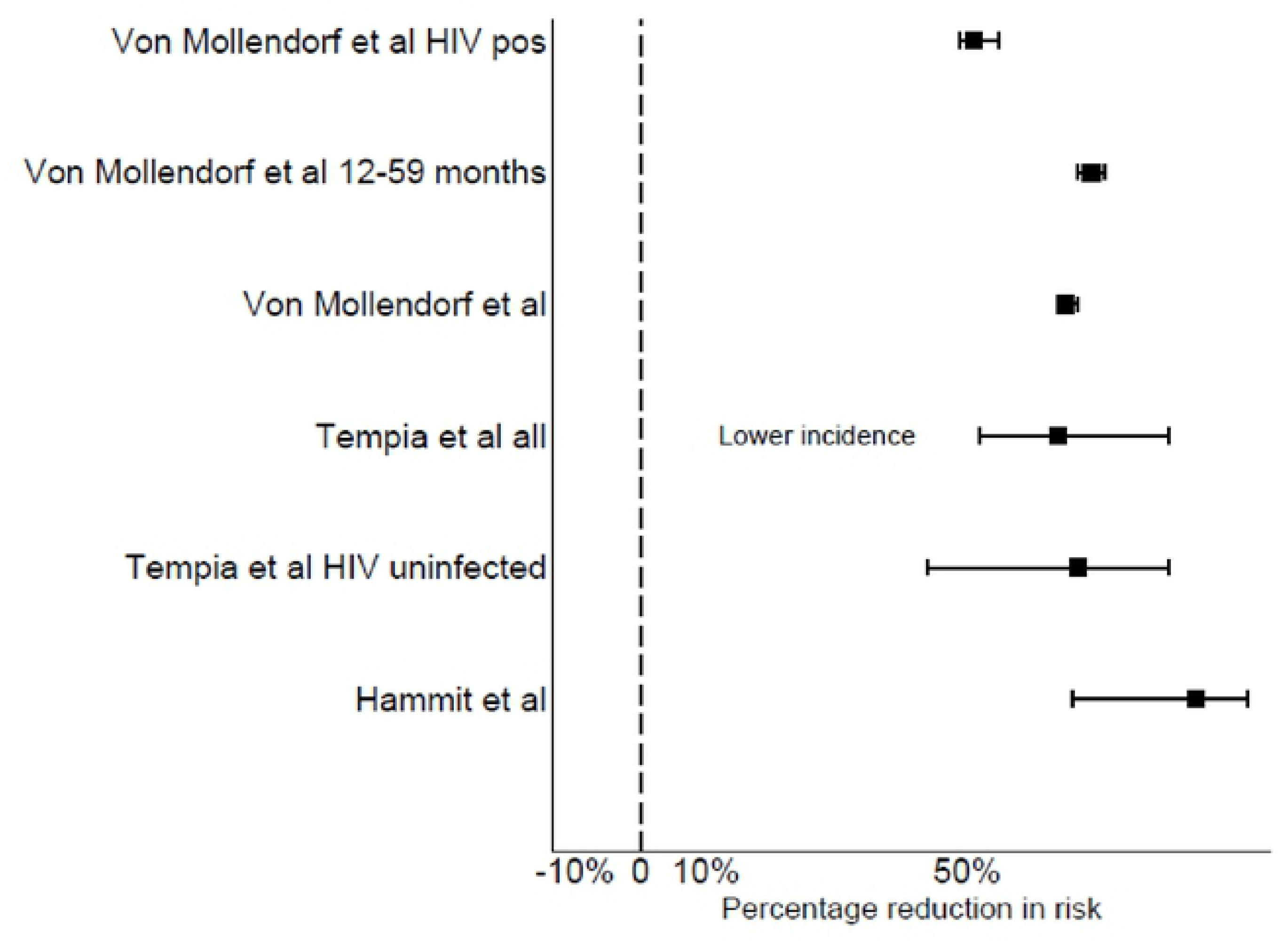
Forest plot showing the percentage risk reduction in pneumococcal pneumonia incidence in included studies and population.

## Discussion

### Summary of evidence

This review set out to answer the question: Does the introduction of pneumococcal conjugate vaccine result in a decline in childhood pneumonia? Since the implementation of pneumococcal vaccination, few studies have evaluated the impact on pneumonia as a specific clinical outcome. Much of the available information on pneumococcal vaccine impact in Africa has been from two countries; The Gambia and South Africa.

There was variability observed in the diagnosis of two of the major outcomes we set out to evaluate; clinical and radiological pneumonia. Only two impact studies consistently applied standardised methods in classifying pneumonia cases(23,30). From our review, we can summarise that the population impact of pneumococcal vaccination on pneumonia depends largely on how pneumonia is classified as an outcome. We observed overall that when the outcome is clinical pneumonia, the impact tends to be modest at best. We see this in the reports by Mackenzie et al from The Gambia, Silaba et al from Kenya and Solomon et al from South Africa. It is important to note however that the severity of pneumonia differed in these studies. Mackenzie and colleagues evaluated pneumonia at the population level and produced results that were not statistically significant in all age groups. Silaba et al however looked at clinical pneumonia hospitalizations (i.e. severe or very severe pneumonia) and showed significant difference in incidence after vaccine introduction. We also see a subtle decline in hospitalizations for clinical pneumonia incidence based on ICD-10 coding. Clinical pneumonia is a non-specific outcome and therefore may not be ideal for evaluating vaccine impact. There was a more positive but still modest impact observed with radiological pneumonia as an outcome. This is likely due to the fact that radiological pneumonia is more specific for pneumococcal disease. Overall, there was a consistent decline in pneumococcal pneumonia cases in the post-vaccine period in reported impact studies(27,36,42). Two studies that reported on vaccine effect in HIV infected children showed that pneumococcal vaccine had similar impact as with HIV uninfected children(36,41).

The overall trend in the findings from this review is similar to reports from other parts of the world. Meta-analyses from other regions have found modest decline in clinical and radiological pneumonia hospitalizations. A meta-analysis using global data from 12 pre-post and time series studies produced a pooled reduction in hospitalization rates for clinical pneumonia of 17% and 9% in the under 24 months and 24 – 59 months’ age groups. However, they reported a more significant decline in hospitalization rates due to radiological pneumonia of 31% and 24% in the same age groups(48). Another meta-analysis of invasive pneumococcal disease hospitalization in Latin America showed a pooled vaccine effectiveness of 8.8% – 37.8% for radiological pneumonia and 7.4% – 20.6% for clinical pneumonia. A randomised placebo-controlled trial on 9-valent vaccine showed vaccine effectiveness of 37% based on per-protocol population(49). Overall, it appears that pneumococcal vaccination has its greatest impact in preventing the more severe forms of pneumonia for which children are likely to be hospitalised. This is probably due to the finding that bacterial pathogens are more likely to cause severe pneumonia(50). The minimal impact on clinical pneumonia might suggest a corresponding increase in other forms of pneumonia or due to serotype replacement(51–53). We observed an age-related trend in vaccine effect with the 12 – 23 months age group experiencing the greatest benefit. This might be due in part to a greater proportion of children under 12 months having not completed their vaccination schedules; and potentially waning immunity in the older age groups(54).

### Limitations

This review has some limitations that have to be considered when interpreting our findings. First was the inconsistency in the definition of pneumonia outcomes between studies, which made combining the impact measured between studies impractical. While some studies used comparable methods for outcome ascertainment, this was not consistent across studies. The WHO standardised definition of clinical and radiological pneumonia is markedly helpful in this situation as it ensures consistency irrespective of study location(55). Studies conducted in locations with a functional health and demographic surveillance system like in the Upper River region of The Gambia and in Kilifi, in Kenya(37,39) were particularly robust as they combined consistent pneumonia surveillance methods with up-to-date population information. Some studied relied on routine clinical data which is usually of variable quality(31,41). One of these studies did not include p-values in its effect measure.

It is also important to note that while we set out to evaluate the impact of 10-valent and 13-valent vaccines, all of the study locations except in Kenya had a period of 7-valent vaccine use. Therefore, is it impossible to separate the effect of the 7-valent from the 10- and 13-valent vaccines since the 7-valent PCV might have changed the pre-existing disease trend.

Like all time-trend studies - including pre-post studies, phenomena such as regression to the mean, seasonality, trend, and history bias have to be considered in the analysis. By including a control outcome and conducting sensitivity analysis, some of the included studies considered the impact of history bias and trend on their results(37). Interrupted time series experiments make adjustments for these phenomena and are a more robust means of assessing vaccine impact(39,56).

Publication bias is one potential limitation of this review; however, we do not believe that it had a significant influence on the findings. This is because we searched multiple databases and search engines to retrieve papers; we also included journals that published African studies and grey-literature. It is therefore fair to assume that the effect of publication bias was minimal in this review. Due to limited resources we were unable to include studies published in other languages; hence, language bias cannot be ruled out in our review.

### Conclusion

To the best of our knowledge this is the first systematic attempt at synthesizing the impact and effectiveness of routine pneumococcal vaccination on childhood pneumonia from this region. The 10- and 13-valent PCV use as part of infant immunization is effective in preventing the more severe forms of childhood pneumonia. There appears to be a smaller effect on clinical pneumonia especially when all severity spectra are included. There is the need for consistency in pneumonia definition for the purpose of disease surveillance and the WHO clinical definitions provide an appropriate option ensuring ease of implementation and reproducibility. One major issue encountered was that few studies had applied comparable pneumonia definitions in estimating disease burden prior to pneumococcal vaccine introduction hence making trend analysis difficult. There is the need for generation of updated information on the causes of pneumonia in this region in this era of extensive vaccine use. Ongoing surveillance is needed to investigate the long-term trend in childhood pneumonia in the post-PCV era in sub-Saharan Africa.

